# Concurrent optogenetic motor mapping of multiple limbs in awake mice reveals cortical organization of coordinated movements

**DOI:** 10.1101/2024.07.05.602302

**Authors:** Nischal Khanal, Jonah Padawer-Curry, Trevor Voss, Kevin Schulte, Annie Bice, Adam Bauer

## Abstract

**Background:** Motor mapping allows for determining the macroscopic organization of motor circuits and corresponding motor movement representations on the cortex. Techniques such as intracortical microstimulation (ICMS) are robust, but can be time consuming and invasive, making them non-ideal for cortex-wide mapping or longitudinal studies. In contrast, optogenetic motor mapping offers a rapid and minimally invasive technique, enabling mapping with high spatiotemporal resolution. However, motor mapping has seen limited use in tracking 3-dimensonal, multi-limb movements in awake animals. This gap has left open questions regarding the underlying organizational principles of motor control of coordinated, ethologically relevant movements involving multiple limbs.

**Objective:** Our first objective was to develop Multi-limb Optogenetic Motor Mapping (MOMM) to concurrently map motor movement representations of multiple limbs with high fidelity in awake mice. Having established MOMM, our next objective was determine whether maps of coordinated and ethologically relevant motor output were topographically organized on the cortex.

**Methods:** We combine optogenetic stimulation with a deep learning driven pose-estimation toolbox, DeepLabCut (DLC), and 3-dimentional triangulation to concurrently map motor movements of multiple limbs in awake mice.

**Results:** MOMM consistently revealed cortical topographies for all mapped features within and across mice. Many motor maps overlapped and were topographically similar. Several motor movement representations extended beyond cytoarchitecturally defined somatomotor cortex. Finer articulations of the forepaw resided within gross motor movement representations of the forelimb. Moreover, many cortical sites exhibited concurrent limb coactivation when photostimulated, prompting the identification of several cortical regions harboring coordinated and ethologically relevant movements.

**Conclusions:** The cortex appears to be topographically organized by motor programs, which are responsible for coordinated, multi-limbed, and behavioral-like movements.

## Introduction

Brain stimulation techniques, (*e.g.*, intracortical microstimulation (ICMS) or optogenetic mapping) have facilitated mapping motor circuitry and provided fundamental discoveries about distributed cortical motor control and common organizational features of the motor system. Penfield and colleagues pioneered electrical stimulation mapping of the human cerebral cortex and revealed the motor and sensory homunculi ([1], [2]). Similar mapping studies have also revealed the cortical organization of monkeys ([3], [4], [5], [6], [7]), squirrels [8], tree shrews [9] and rats ([10], [11]). These studies reveal that mapping motor movements beyond cytoarchitecturally defined motor cortex (*e.g.*, primary (M1) and secondary (M2) motor regions) is crucial for a comprehensive understanding of action generation and planning. For example, motor movements can be elicited from somatosensory regions in multiple species [11]. In rodents, forelimb movements are controlled by two distinct cortical functional areas: a smaller rostral forelimb area (RFA) in M1, and a larger caudal forelimb area (CFA) overlapping with portions of M1, M2 and primary somatosensory (S1) cortex. More recent work has identified that motor cortex contains functional zones which integrate inputs from multiple limbs and orchestrate complex motor programs [12]. Further supporting this idea is evidence showing that motor cortex and anterior and posterior parietal cortex form a sensorimotor integration network [13]. These and other observations beg the question of whether more “behavior-like” actions are spatially encoded across the entire cortex.

Understanding this multi-limb integration requires moving beyond the conventional somatotopic body map. Recent research highlights the presence of an additional organizing principle known as the action map, where functional zones associate with ethologically relevant movements or “actions” [14]. Furthermore, fMRI methods suggest that M1 interweaves two parallel systems: effector-specific (foot, hand, and mouth) regions for isolating finer movements and a system for whole-body action planning [15]. Using ICMS, action maps have been demonstrated in monkeys ([4], [16], [17]) and even to a limited extent, in humans [18], and include actions such as reaching, hand-to-mouth, grasping, and defensive movements [16]. Similarly, distinct functional zones within the rodent M1 have been identified. Selective inactivation of the RFA in rats during skilled forelimb behaviors indicates a movement-based functional organization of the motor cortex [10], a perspective supported using long-train (LT) ICMS [19]. In mice, LT optogenetic stimulation of the motor cortex evokes abduction and adduction of the forepaw [20], along with discrete and rhythmic forelimb movements [21], highlighting functional subregions within the forelimb cortex for different motor dynamics. Recently, LT-ICMS has been used to evoke complex movements (*e.g.*, limb elevation, retraction, and “digging”) from the historically defined CFA and RFA, resembling movements observed during natural behavior [22]. However, a comprehensive characterization of complex movements simultaneously including forelimbs, hindlimbs, and mouth in mice has not yet been documented. Furthermore, cortical stimulation has primarily focused on somatomotor regions and has not encompassed whole-hemisphere interrogation, despite the potential presence of regions implicated in ethologically significant movements.

Here, we integrate optogenetic stimulation with DeepLabCut (DLC) [23], a deep learning driven pose-estimation toolbox, to develop Multilimb Optogenetic Motor Mapping (MOMM), enabling precise localization of coordinated movements across forelimbs, hindlimbs, and mouth in three dimensions. We validate the efficacy of MOMM by quantifying directional translations of each limb and demonstrating consistent cortical motor representations across multiple trials and subjects. These representations extend beyond the cytoarchitecturally defined somatomotor cortex, revealing that 64% of the cortex, including 14% outside traditional sensorimotor areas, contributes to motor output upon photostimulation. MOMM also discerns subtle articulations of the forepaws that reside within motor representations of the forlimb. Moreover, many cortical sites exhibit concurrent limb coactivation when photostimulated, and harbor coordinated and ethologically relevant movements.

## Methods

### Experimental Summary

We developed a Multilimb Optogenetic Motor Mapping (MOMM) method with the goal of understanding cortical organization and execution of multi-limbed movements in awake mice. Optogenetic photostimulation was integrated with multiple feature tracking using DeepLabCut (DLC) in order to map behaviors on the cortex. Mice were extensively acclimated to awake MOMM prior to mapping. 378 cortical sites covering the convexity of left hemisphere were photostimulated using a 473 nm laser and evoked movements were recorded using four CMOS cameras positioned around the animal. DLC was used to track features of interest and pairs of cameras were triangulated into front (forelimbs and mouth) and hind (hindlimbs) views for characterizing complex motor output. Gross translations were quantified using the area under the total displacement curve within the stimulus window and mapped onto the cortex. Some movements (*e.g.*, forelimbs) were further analyzed to determine the topography of finer articulations (*e.g.*, forepaw rotations). Investigation of sites co-activating multiple limbs identified the presence of cortical motor programs which give rise to complex, coordinated, and multi-limbed motor movements.

### Mice

Nine Thy1-ChR2-YFP mice (3 females, 6 males) [24], aged between 3.5 and 7 months were used for experimentation. The spatial distribution of ChR2 in this mouse line has been well characterized in previous studies ([24], [25]) and is primarily expressed within the axons and dendrites of the layer V pyramidal neurons. These neurons have pronounced apical dendritic tufts in layers 1 and 2/3 ([26], [27], [28]). Mice were given *ad libitum* access to food and water with a 12 hours on – 12 hours off light cycle. All experimental protocols were approved by the Animal Studies Committee at Washington University in Saint Louis.

### Animal Preparation

Prior to motor mapping, each mouse’s skull was fit with a small Plexiglas window as previously described ([29], [25]). In brief, a single injection of buprenorphine-SR (1.0 mg/kg subcutaneous) was administered 1 hour before surgery. Mice were then anesthetized with isoflurane (3% induction, 1% maintenance, 0.5L/minute air) and allowed 2-5 minutes for anesthetic transition. Mice were maintained at 37°C through rectal feedback and a heating pad (mTCII, Cell Microcontrols, Norfolk, VA, USA). A midline incision was made along the top of the head and the scalp was retracted, keeping the skull intact. A Plexiglas window with pre-tapped holes, sterilized with Chlorhexidine, was installed over the skull, and secured with dental cement (C&B-Metabond, Parkell Inc., Edgewood, NY, USA). After surgery, the mice were returned to their cages and monitored for seven days, during which time no handling or optogenetic stimulation was performed.

### Motor Mapping Stand and Acclimation

The system used for motor mapping is a modified version of a previously published system used for mapping cell-specific effective connectivity in the mouse brain (**Fig. 1A-B**) [29]. A custom motor mapping head-fixed mounting system was designed, and 3D printed (print files available on request) to facilitate unrestricted, naturalistic, movement of all limbs and mouth. Mice were suspended at the trunk using a Velcro strip attached to the 3D printed mount and were head-fixed in a stereotactic frame using the cranial window, as per prior reports ([30], [29], [25], [31]). A partition was placed between the fore- and hindlimbs to prevent the limbs from being captured in opposed camera views (see below) and confusing DLC tracking but did not restrict movement.

**Figure 1.**
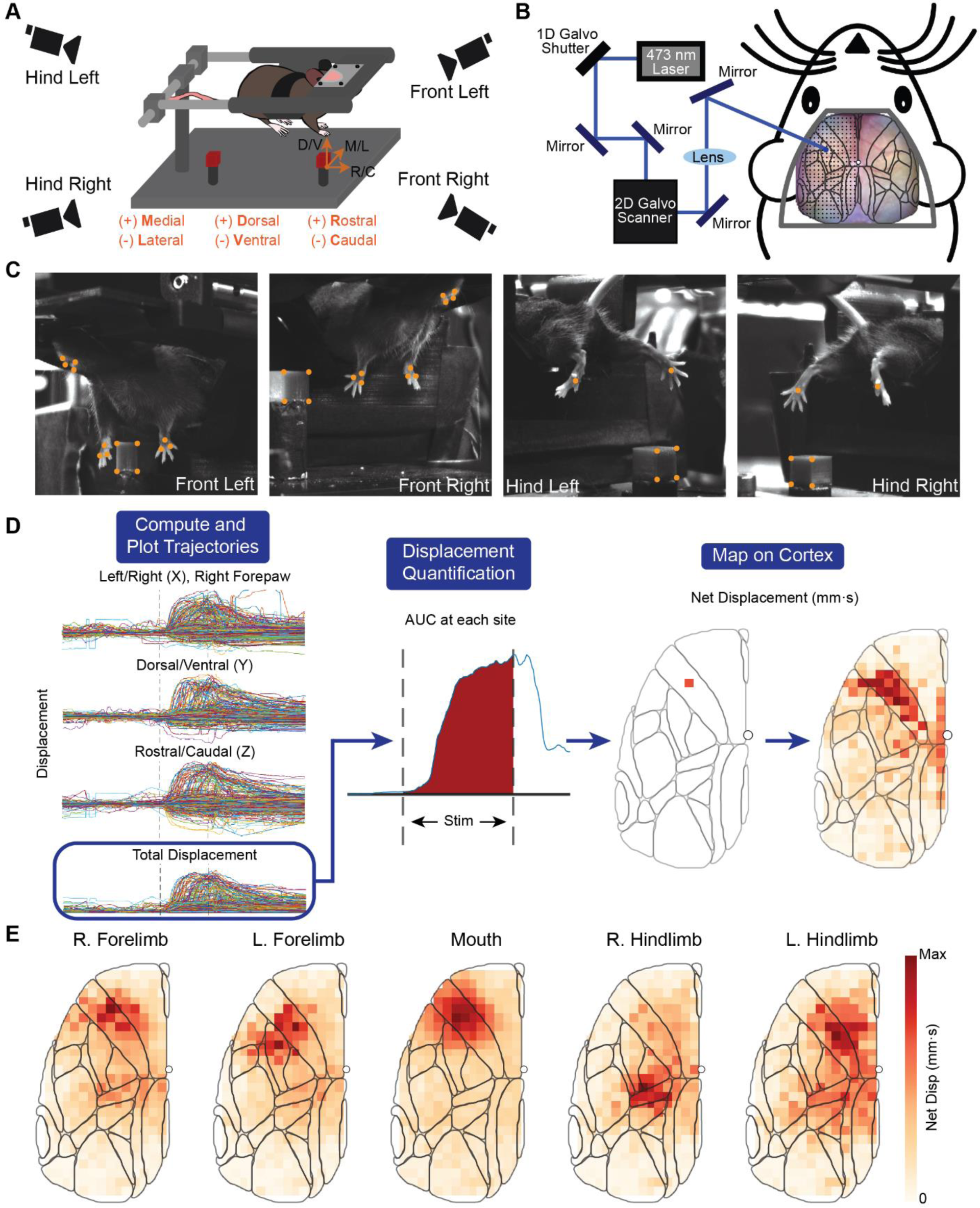
MOMM hardware and analysis. A) Schematic of the motor mapping stand and feature-tracking cameras. Awake, behaviorally acclimated mice are head fixed and suspended with all four limbs freely hanging to facilitate unrestricted, naturalistic movements. Four cameras are positioned around the mouse, as shown. Movement trajectories are registered into a common space via a rigid transform to a permanent cube present in all camera views and sessions; basis vectors of this space are indicated as dorsal/ventral, medial/lateral, and rostral/caudal directions. B) Simplified schematic of the laser guiding system for photostimulation. A 473 nm laser was guided using a series of mirrors and galvos (left) to stimulate a cortical grid of 378 sites (300 micron separation) covering most of the left hemisphere (right). The stimulation grid was registered to the mouse cortex using an affine transform prior to scanning. The white dot indicates bregma. C) Field-of-views of the four CMOS cameras, each recording at 200 Hz. Features labeled and tracked by DeepLabCut, and used in movement quantification are indicated in orange. Pairs of views were used to triangulate trajectories into 3D (Front Left/Right for forelimbs and mouth, Hind Left/Right for hindlimbs). D) Motor Mapping Pipeline: an overview of the pipeline through which behavioral videos are converted into movement trajectories and mapped onto the cortex as displacement. In brief, DLC models were trained to label and track features in each view. Features were triangulated and co-registered prior to plotting trajectories along each direction. The Euclidean norm was computed at each time point to generate the total displacement trace, and net displacement was quantified as the area under the curve (AUC) of this trace at each site. This value was plotted onto the cortex for all sites to generate a motor map and quantify gross displacement of each limb and mouth. E) Motor maps quantifying the gross displacement of each limb and mouth. The net displacement value at each stimulation site is the median across the group. The limbs displayed are (left to right) Right Forepaw (Peak: 9.86 mm•s), Left Forepaw (Peak: 10.9 mm•s), Mouth (Peak: 0.644 mm•s), Right Hindpaw (Peak: 4.10 mm•s), Left Hindpaw (Peak: 3.02 mm•s).

A major advantage of optogenetic stimulation is the ability to perform mapping in awake mice. This necessitates rigorous behavior acclimation to the motor mapping stand as spontaneous (non-photo-evoked) movements may obscure stimulus-evoked movements. To this end, mice were trained over a 5-day period using a modified version of previously published protocols ([29], [31], [32], [33]). During the first 2 days, mice were accustomed to handling for 30 minutes/day. The following 2 days, mice were restrained within the mounting system for 1 hour with no stimulation. Finally, on the 5^th^ day, mice were acclimated to optogenetic stimulation by stimulating each site a single time.

### Optogenetic Hardware and Data Acquisition

A EMCCD camera (Andor iXon 897, Oxford Instruments, Oxfordshire, England) was positioned above the mouse and used to register the laser stimulation sites to the cortical surface. This registration was achieved through an affine transformation using user-defined selections of the junction between the coronal suture and sagittal suture (anterior) and lambdoid suture (posterior) as landmarks, and a third constructed point, which ensured no shear. A 473 nm laser (BL473T-150, Shanghai Laser & Optics Century Company, Shanghai, China) with a beam diameter of approximately 50 microns was optically shuttered using a 1-D galvanometer (Thorlabs GVS002), reflected off 2 broadband mirrors (Thorlabs BB1-E02), and delivered to the cortex by 2-D galvanometer (Thorlabs GVS002) scanning mirrors and periscope optics (**Fig. 1B**).

To evoke motor movements, the photostimulus design followed previous work ([26], [20], [21]) and consisted of 2.5 second blocks, with a 500 ms stimulation train (10 ms pulses delivered at 50 Hz) with a laser power of 5 mW. One second was provide before and after each stimulus to promote a return to resting posture and ensure full execution of photo-elicited movement programs. Photostimulation was performed over a grid encompassing the left hemisphere and consisting of 378 sites separated by 300 microns (**Fig. 1B**). Before experimentation, the order of photostimulation sites were randomized, with a constraint that directly neighboring (300-micron radius) sites were not stimulated in sequence. This randomized sequence was used for all mice. Motor movements were recorded at 200 Hz by four cMOS cameras (Allied Vision Mako U-130B USB 3.0, Monochrome Camera) positioned around the mouse and separated by approximately 45 degrees (**Fig. 1A**). To ensure frame synchronization between the four cameras, cameras were hardware triggered *via* a JST 7-pin cable (Allied Vision #14-155) connected to a data acquisition card (DAC) (PCI 6733, National Instruments, TX, USA). Movements of the right and left forelimbs and mouth were captured by the two front cameras (Front-Left, Front-Right) while movements of the right and left hindlimbs were captured by the two rear cameras (Hind-Left, Hind-Right) (**Fig. 1C**). The fields-of-view (height x width) of each camera were approximately: Front-Right: 69×69mm (11.6 pixels/mm); Front-Left: 72×72mm (11.1 pixels/mm); Hind-Right: 75×75mm (6.5 pixels/mm); and Hind-Left: 69×69mm (11.6 pixels/mm). The fields-of-view of the two pairs of tracking cameras were triangulated using the DeepLabCut-3D pipeline (see below). To co-registration movement trajectories from each camera-pair’s, 3D-space (*e.g.*, Front), a reference cube of known side length (1 cm) was placed within all FOVs. Movements were quantified along three axes: rostral/caudal, dorsal/ventral, and left/right (**Fig. 1A**).

To ensure synchronization between both the stimulation and camera recordings, trigger control of the photostimulation laser and cameras were handled by the DAC. Owing to differences in cranial window placement, not all sites coincided within the cortex for each mouse. For these mice, sites outside the exposed cortex were excluded from stimulation and analysis. Thus, awake mice were mapped between 900 and 960 seconds depending on the number of sites registered within the exposed cortex for each mouse. 10 consecutive motor mapping sessions were performed on each mouse.

### Data Processing

All DeepLabCut processing was performed through Anaconda on Windows 10 using a NVIDIA Quadro A5000 GPU. Subsequent analysis was performed using MATLAB 2022b and 2023a (Mathworks).

### *3-Dimensional* Pose Estimation and Movement Quantification

#### DeepLabCut

DLC’s markerless pose estimation toolbox was used to label and track features in each camera view (shown in orange, **Fig. 1C**). Specifically, features on the right and left forelimbs, right and left hindlimbs, and mouth were labeled. Training was performed on a random subset of our data (∼10%) using DLC 2.2 with default network parameters. To facilitate three-dimensional pose estimation, the fields-of-view of all individual tracking cameras were triangulated using the DLC-3D pipeline [34]. The Front-Right and Front-Left views were triangulated for forelimbs and mouth movements while Hind-Right and Hind-Left were triangulated for hindlimb movements. The camera-facing corners of the cube present in each camera view (front face for Front views, rear face for Hind views) were tracked and labeled by DLC, and the lower left corner was defined as the origin. Owing to slight errors in DLC labeling and DLC-3D triangulation, the four corners may not always be coplanar and separated by exactly 1 cm. To address this concern, the most orthonormal set of adjacent edges was identified by calculating the dot product between all adjacent edges. The pair of edges with the smallest dot product (*i.e.*, the most orthogonal) were then used to perform a Gram–Schmidt orthonormalization (which forces orthonormality) to generate a new set of basis vectors spanning the 2D space in the medial/lateral and dorsal/ventral plane. The last basis vector (rostral/caudal) was calculated from the cross product between of the previously derived basis vectors. Finally, a rigid transformation was performed from the native DLC-3D basis vectors to the new cube-derived basis vectors.

### Gross Limb Displacement

For the fore- and hindlimbs, the changes in position of the center feature of each paw (center of the metacarpal) was used to quantify the translation for the associated limb. For the mouth, this translation was calculated as temporal-variations in the Euclidean distance between the upper and lower lip. Trajectories at each site were plotted as directional displacements to visualize movement along each axis (**Fig. 1D**, panel 1). Each movement trajectory was baseline subtracting using the mean of the first 0.5 seconds (t = −1.0s to t = −0.5s) of the block. To compress these directional displacements, total displacement was computed as the Euclidean norm of directional displacements at each time point (i.e., 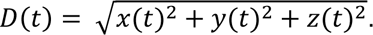 Next, the area under the curve (AUC) of the total (gross) displacement trace during the stimulus window (t = 0s to t = 0.5s) was calculated (**Fig. 1D**, panel 2) and mapped to the corresponding cortical location. This was repeated for all sites to generate a motor map (**Fig. 1D**, panel 3). This process was repeated for all four limbs and the mouth across all trials of each animal. The median total displacement at each stimulation site across the group was mapped to generate group level gross translation motor maps (**Fig. 1E**).

### Directional Limb Movements

For the forelimbs and hindlimbs, the gross translations were also parsed into their directional components and mapped (**Fig. 2A**, **Sup. 2A**). As in the gross translation case, the medial/lateral, dorsal/ventral, and rostral/caudal trajectories were integrated over the stimulus window and these displacement values were plotted onto the cortex at the corresponding stimulation site. For the left limbs, the displacements along the medial/lateral direction were inverted to ensure that medial limb movements corresponded to positive displacements.

**Figure 2.**
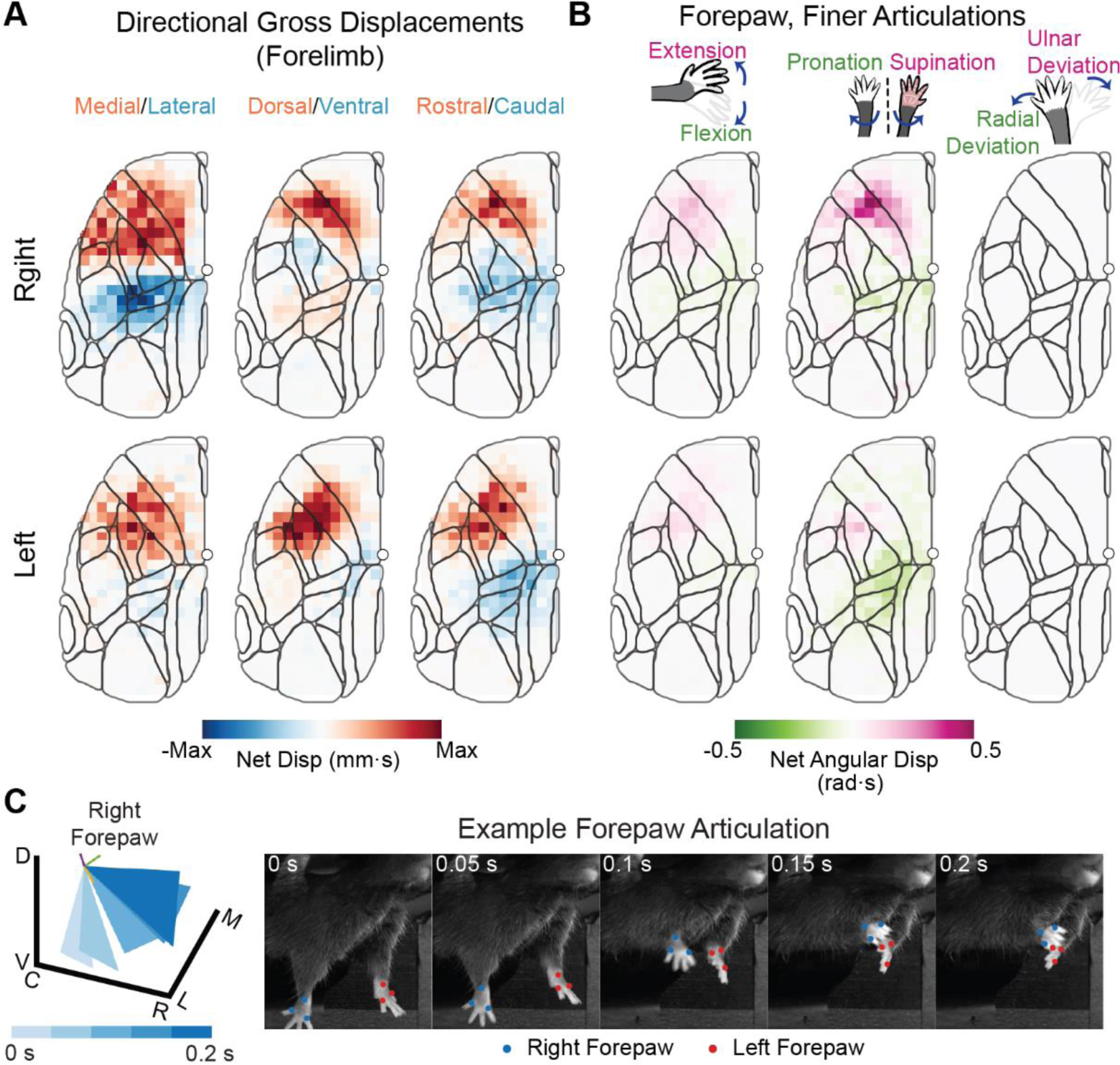
Gross forepaw movements can be dissected into directional displacements and finer, forepaw articulations. A) Motor maps showing the directionality of right (top) and left (bottom) forelimb movements along three directions. Displacement was mapped using AUC values as in the case of gross displacements but using the directional trajectories shown in Fig. 1D. Positive (red) values indicate movement in the medial, dorsal or rostral directions while negative (blue) values indicate movement in the lateral, ventral or caudal directions. Maximum displacements of right forelimb: 2.39 mm•s (M/L), 4.85 mm•s (D/V), 8.04 mm•s (R/C), left forelimb: 4.91 mm•s (M/L), 4.45 mm•s (D/V), 8.57 mm•s (R/C). B) Three DLC-labled points on the forepaw (see Fig. 1B) were used to construct a plane. The gross displacement was subtracted from these points and the Euler angles were computed using this plane to quantify three rotations: extension/flexion, pronation/supination, radial/ulner deviation. The primary rotations of both forepaws over M1 are supination movements (peak: supination, 0.505 rad*s, area: 8.66 mm^2), followed by extension (peak: extension 0.198 rad*s, area: 3.47 mm^2). There is a negligible amount of radial/ulnar deviation within both forepaw movements. C) The position of the forepaw plane is plotted in three dimensions (left), with each time point corresponding to a movie frame from an exemplary trial (right).

### Finer Forepaw Articulations

Often, forelimb movements exhibited more articulated movements (*e.g.*, forepaw rotations) during gross forelimb motions. To quantify these finer movements, a plane was constructed using three features placed on the forepaw: the wrist (origin), proximal knuckle and distal knuckle (**Fig. 1C**). The origin (*i.e.*, the wrist) was subtracted from these three points to remove gross translation and to isolate rotations about the wrist. A 3^rd^-order lowpass Butterworth filter with a cutoff frequency of 20 Hz was applied to the trajectories at each point. Rotational axes were defined by 1) the normal line at the center of the plane, 2) a line extending from the origin that bisected the other two points, and 3) the cross product between these two lines (**Fig. 2C**, left). Euler angles were then computed to quantity the three rotations: extension/flexion (pitch), pronation/supination (roll) and radial/ulnar deviation (yaw). These rotational angles were computed at each timepoint for both forepaws and, as in the directional limb movement case, the AUC of each rotational trajectory during the stimulus period was mapped onto the cortex (**Fig. 2B**). To help visualize the rotational movements, the position of the forepaw plane was plotted in three dimensions (**Fig. 2C**, left), with each time point corresponding to a movie frame from an exemplary trial (**Fig. 2C**, right). An exemplary trial supplemental video is also provided.

### Manual Scoring of Motor Movements

To establish a definitive benchmark for all evoked limb movements, motor responses from a single trial per mouse were manually categorized. The movements of all limbs and mouth at each stimulation site were categorized utilizing the subsequent descriptors: right/left forelimb extension, right/left forelimb retraction, right/left hindlimb flexion, mouth opening, right forepaw grasping, or absence of movement. The movements of the forelimbs were further categorized as discrete or rhythmic. The incidence of each movement class was then mapped on the cortex, with the numerical value at each site denoting the count of mice exhibiting that specific movement pattern (**Fig. 3**).

**Figure 3.**
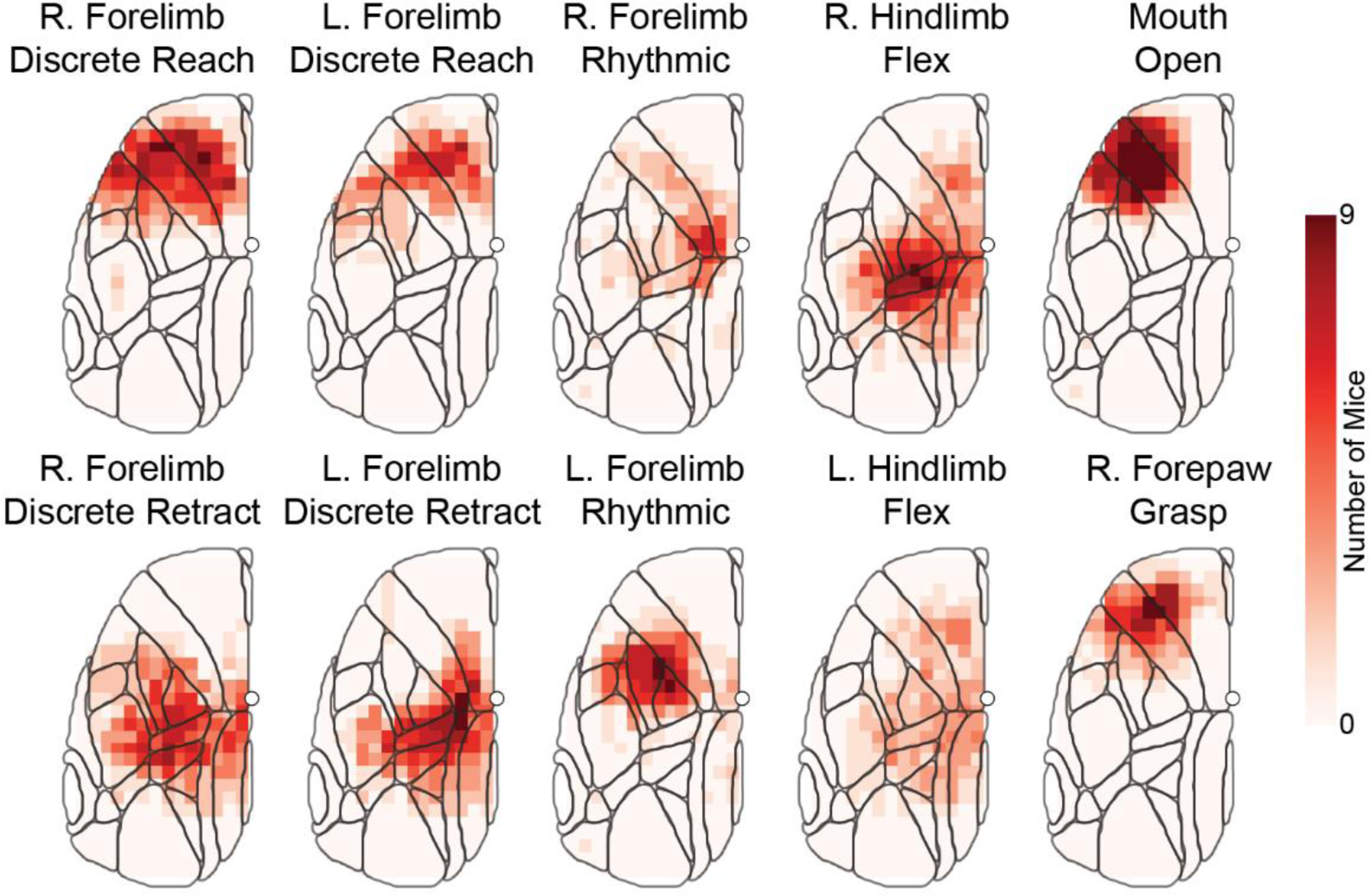
Coordinated movements are corroborated by empirical analysis of evoked movements in individual mice. Motor responses from a single trial of each mouse were manually assessed and the movements of all limbs and mouth at each stimulation site were categorized utilizing the labeled descriptors. Movements of the forelimbs were further categorized as discrete or rhythmic. The numerical value at each site denotes the count of mice exhibiting that specific movement pattern.

### Assess Regions of Overlap

The gross translation maps for each limb exhibit a bimodal distribution, demonstrating topographical segregation about bregma. To delineate these clusters of movements, the motor maps were partitioned into regions above and below bregma. After this division, the maps were thresholded, isolating the top 33rd percentile of activity, and spatially smoothed using a 9×9 gaussian kernel (2 pixel standard deviation). Subsequently, contours were delineated around the resulting clusters. The partition bearing the greater maximum displacement was designated as the “primary” representation for that limb, while the other partition was labeled as “secondary”. The mouth motor map displayed a unimodal distribution, resulting in a single contour for that case. These contours were then superimposed to evaluate the extent of overlapping stimulation sites across different limbs. (**Fig. 4A**, **4D**).

**Figure 4.**
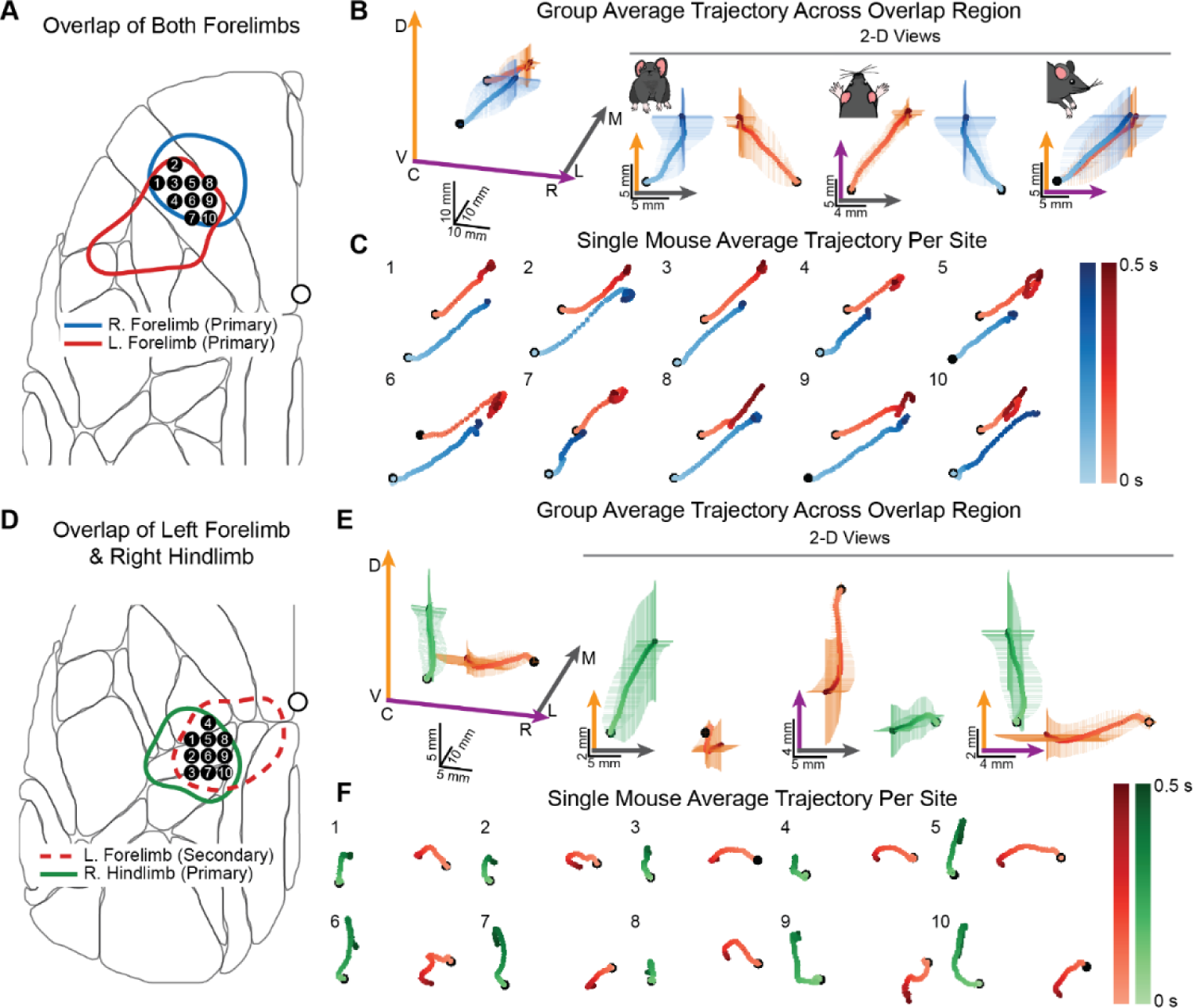
Coordinated and multi-limbed movements reside within the overlap of gross displacements. Contours were delineated around the top 33^rd^ percentile of evoked motor movements for each limb. (A) Intersection of contours of the primary representations of the right and left forepaw motor maps. The ten stimulation sites located within the overlap region are indicated with black circles. (B) The motor movement trajectory at sites within the region of overlap were averaged across all runs and mice to generate the group average trajectory. The 2D viewpoints of the group average trajectory are provided to enhance interpretability of the evoked movement. Only movement during the stimulus period is displayed and the trajectories are color coded in time with darker colors indicating the end of the stimulus train. (C) Limb movement trajectories from a representative awake mouse following photostimulation of the 10 overlapping sites. Each trajectory is the average across all runs of mouse at the indicated stimulation site. (D-F) Same as A-C but within sites of overlap of the secondary left forelimb and primary right hindlimb contours.

The motor movement trajectory at sites within the region of overlap were averaged across runs and mice to generate the group average trajectory (**Fig. 4B**, **4E**). The viewpoint of all 3D trajectories was angled from the front right of the animal (azimuth: 105 deg, elevation: 25 deg in MATLAB). The group average trajectories were projected onto 2-dimensional planes to enhance interpretability of movement patterns. These trajectories were color-coded across time, with lighter shades denoting the onset of stimulus and darker shades signifying its termination.

Standard error bars were incorporated at each timepoint along each directional axis. To assess the consistency of these movements at the individual mouse level, trajectories from a single mice were averaged across all trails and plotted for each stimulation site within the overlapping region (**Fig. 4C**, **4F**). The degree of consistency was quantified for all nine mice by correlating site level trajectories with the group average trajectory spanning the entire overlap region. The average correlation and the standard deviation across all sites of overlap are reported (**Table S1**).

To quantify the spatial specificity of overlapping movements, covariance was computed between the group average trajectory at sites surrounding the region of overlap and within the region. This analysis was performed along the anterior-posterior and medial-lateral directions, separately for each limb. For this analysis, before covariance calculating, 3-dimensional trajectories were flattened into single-column vectors. The covariance was then plotted as a function of cortical position, highlighting sites corresponding to the overlap region. (**Fig. S3**).

### Action Mapping

Movements within the overlapping zones of multiple limbs unveiled ethologically relevant behaviors. These contours were overlayed onto a single map, where solid lines represented “primary” movements and dashed lines indicated “secondary” movements (**Fig. 5A**). The convergence of multiple contours was emphasized with a darker contour, each delineated with ethologically meaningful descriptors (actions). Frames from exemplary mouse movements were presented for each action, showcasing specific behaviorally relevant movements (**Fig. 5B**). In cases involving both fore and hindlimbs, frames from two viewpoints of the mouse were included (**Fig. 5C**).

**Figure 5.**
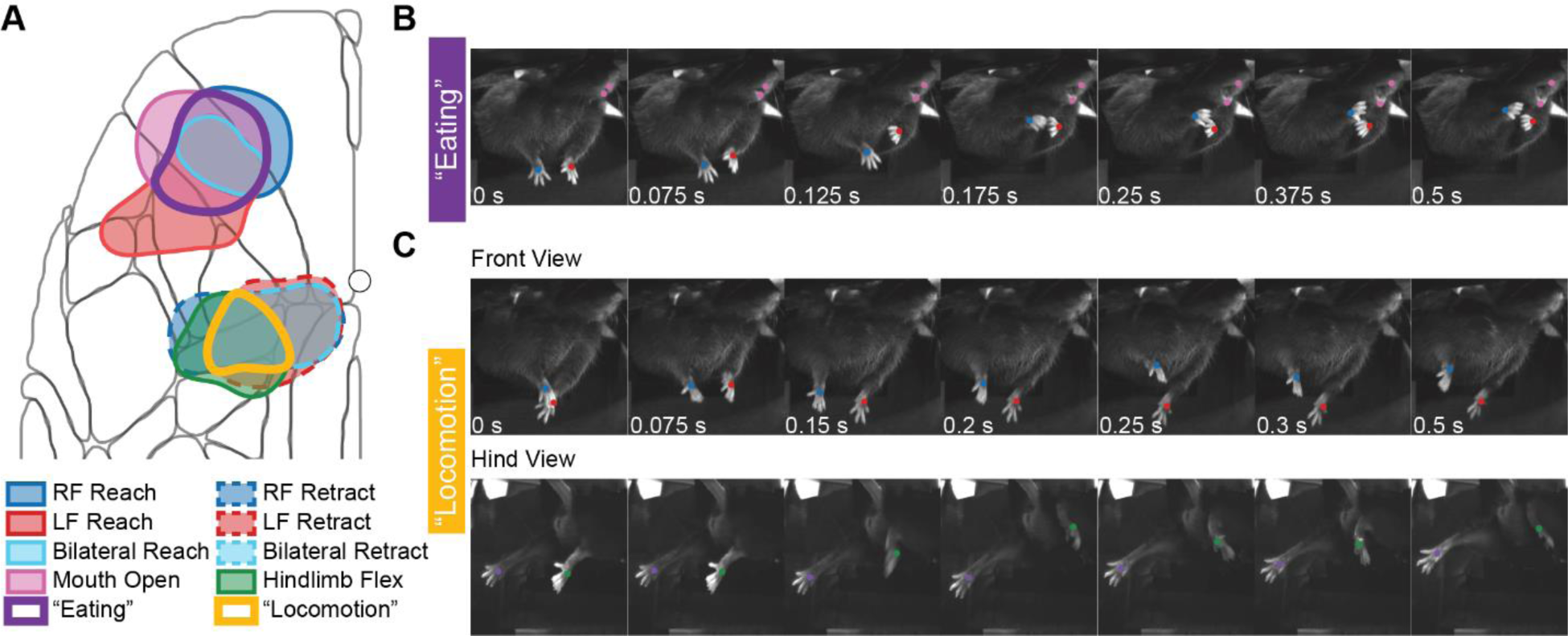
Coordinated, multi-limbed, movements are spatially organized on the cortex. A) Contours of all limbs and mouth are overlayed on the cortex. Solid lines represent primary movements and dashed lines indicate secondary movements. The convergence of multiple contours is indicated with a heavier contour, and each delineate an ethologically relevant movement. B) Image sequence of example mouse displaying an “eating” action where both forelimbs reach towards the mouth concurrently as the mouth opens. C) Image sequence of example mouse displaying an “locomotion” action characterized by retracting and rhythmic forelimb movement in conjunction with a flexion of the hindlimbs.

### Variability in Motor Mapping

Mouse-level variability in gross translation was computed for each limb and the mouth. Specifically, the area under the curve (AUC) displacement at each site across all runs for each mouse was subjected to t-tests (5% significance level) to assess the significance of evoked movements against the null hypothesis of no movement. Additionally, a median map of t-values was generated using the median t-statistic for each site (**Fig. S1**). This analysis was repeated for directional limb movements, with only the median maps for each direction displayed (**Fig. S2B**).

## Results

### Concurrent mapping of multiple features using MOMM

We used MOMM to establish topographical cortical organization of motor control of the forelimbs, hindlimbs, and mouth (**Fig. 1**). Photoelicited motor movements at each site were evaluated by calculating the area under the curve (AUC) of the evoked movement trajectory during the stimulus presentation and mapping that value to the associated cortical location (**Fig. 1D**). These displacements were ttest (5% significance level) at each site across all runs of each mouse and the topographies of the median ttest maps closely resemble those of the gross translation maps (**Fig. S1**). Cortical movement representations of the right forelimb (**Fig. 1E**, Panel 1) exhibit a rostral representation largely overlapping with primary and secondary motor cortex and somatosensory regions adjacent to a caudal representation within posterior somatosensory regions, and portions of parietal and retrosplenial cortex. Representations of the left forelimb (**Fig. 1E**, Panel 2) exhibit a similar topography as the right forelimb, with more of the rostral representation overlapping with somatosensory forelimb cortex and secondary motor cortex. Directional translation maps for the right (**Fig. 2A**, Top) and left forelimb (**Fig. 2A**, Bottom) each exhibit bimodal distributions separated about bregma corresponding to forward (rostral) (**Movie S1A**) and retracting movements (caudal) of each limb (**Movie S1B**). For both forelimbs, larger photo-evoked movements (greater limb displacement) were observed in the rostral representations.

Mapping of the hindlimbs reveals a different picture. Larger right hindlimb movements (redder) overlapped with somatosensory hindlimb and trunk cortex while smaller movements occurred when stimulating primary and secondary motor cortex (**Fig. 1E**, Panel 4). Conversely, the largest movements of the left hindlimb (**Fig. 1E**, Panel 5) occurred within primary and secondary motor cortex while smaller movements resided within posterior parts of somatosensory cortex and extended to parietal and retrosplenial regions. Interestingly, while motor maps of the forelimbs and hindlimbs each exhibited spatial separation about bregma, the “rostral” and “caudal” portions of the hindlimb maps do not correspond to forwards (rostral) and backwards (caudal) hindlimb movements (**Sup. 2A**). Despite this phenomenological difference, these two spatial distributions will still be referred to as “rostral” and “caudal” to maintain consistency with prior nomenclature used in the literature for the forelimb.

Mapping motor movements of the mouth (**Fig. 1E**, Panel 3) revealed a single, topographically smooth cluster centered over lateral portions of primary motor cortex that extended into primary somatosensory forepaw and mouth. Taken together, these results suggest that motor movement representations extend beyond cytoarchitecturally defined somatomotor cortex and involves higher-order brain regions associated with sensory integration and executive function (parietal) ([35], [36]) as well as those involving cognition, planning, and navigation (retrosplenial) [37].

### Finer articulations of forepaw are spatially superimposed on gross displacement maps

In line with prior work ([20], [10], [21], [22]), forelimb motor maps exhibit distinct rostral and caudal representations. LT-ICMS within these regions results in reach- and grasp-like motions qualitatively described in rats ([38], [10], [11]), and mice [22]. Taking advantage of the high-fidelity tracking afforded by DLC, three features on the paw were used to define a plane (**Fig. 1C**), namely the wrist (origin), proximal knuckle and distal knuckle. For both forepaws, Euler angles were computed at each timepoint to quantity the three rotations: extension/flexion (pitch), pronation/supination (roll) and radial/ulnar deviation (yaw). These rotations mapped to each photostimulated site by calculating AUC of each rotational trajectory during the stimulus period. Visualization of these maps (**Fig. 2B**) demonstrates that finer forepaw articulations are also spatially localized on the cortex and confined to regions corresponding to gross right forepaw movements. In fact, these forepaw articulations superimpose upon the gross forelimb displacements, indicating there is often co-activations of both gross and finer movements. Each rotational movement resides primarily over the rostral representations of both forelimbs. For the right forepaw, the rotational movements are primarily composed of a forepaw extension (peak: 0.198 rad·s) combined with a supination (peak: 0.505 rad·s) with a negligible amount of radial/ulnar deviation (<<0.01 rad·s). The left forelimb rotations, though smaller in magnitude, also compose of an extension (peak: 0.107 rad·s) combined with a supination (peak: 0.184 rad·s) with negligible radial/ulnar deviation (<<0.01 rad·s). For both paws, the combination of these rotational movements produces grasping-like motions (**Fig. 2C**, **Movie S2**).

### Coordinated and multi-limbed movements reside within the overlap of gross displacements

The directional translation maps (**Figs. 2A**, **S2A**) revealed significant variations in motor movements across all three axes and the four limbs. To establish a baseline range of limb motions, motor responses from a single trial per mouse were manually categorized. Movements at each stimulation site included: right/left forelimb extension, right/left forelimb retraction, right/left hindlimb flexion, mouth opening, right forepaw grasping, or no movement. Forelimb movements were further classified as discrete or rhythmic. Mapping the incidence of each movement class on the cortex revealed topographies reflecting gross limb displacements (**Fig. 3**). Notably, manual scoring revealed an overlap between discrete right- and left-forelimb reach and mouth opening, aligning with the spatial coincidence of forepaw rotations (**Fig. 2B**) with maps of gross forepaw and mouth displacement (**Fig. 1E**). In fact, most of the manually scored maps overlap to some degree, a finding mirrored by the gross limb displacements, indicating that photostimulation of a single site can evoke movements simultaneously across multiple limbs.

We hypothesized that these sites of overlap correspond to specific cortical regions subserving coordinated motor output. To test this hypothesis, we first generated a set of contours by thresholding each motor map (mouth and separately for rostral/caudal representations) to only consider the top 33% of motor displacements. Photoevoked movements from sites within overlapping contours were then investigated for the presence of co-activated limb movements (**Fig. 4**). For example, photostimulation of any of the ten sites residing within the intersect between the rostral contours of the right and left forepaw (**Fig. 4A**) results in bilateral reaching motions executed by both forelimbs (**Fig. 4B**, **Movie S4A**). This observation was highly consistent across all sites within the region of overlap, even at the single mouse level (**Fig. 4C**) and was assessed through correlations between the group average and individual, mouse-level, site trajectories (**Table S1**). To determine the spatial specificity of this bilateral movement, photoevoked movements at sites along the anterior/posterior and medial/lateral axes of the region of overlap were quantitively compared with the group-averaged movement within the overlapping contours (**Fig. S3)**. Along the anterior/posterior axis, site-wise movement trajectory covariance increased at closer proximity to the overlap region for both limbs and reached a maximum within the contoured region (Posterior RF: −0.793±10.0, Peak RF within overlap: 36.9, Anterior RF: 25.0±13.8, Posterior LF: 11.9±15.1, Peak LF within overlap: 55.1, Anterior LF:19.1±11.5). Along the medial/lateral axis, a similar trend was observed for the left forepaw, while movements of the right forepaw covaried similarly with movements observed across all sites investigated (Medial RF: 20.6±12.2, Peak RF within overlap: 36.9, Lateral RF:19.5±9.90, Medial LF: 15.0±14.9, Peak LF within overlap: 55.1, Lateral LF:26.7±17.5). These findings suggest a sharp topographical gradient for this coordinated movement.

To determine whether these observations extended to other cortical sites of overlap, we also investigated the intersection between the caudal representations of left forelimb and right hindlimb (**Fig. 4D**). Photostimulation of any of the 11 overlapping stimulation sites resulted in left forelimb retraction juxtaposed with a large cyclic motion in the right hindlimb (**Fig. 4E**, **Movie S4B**). As above, this retraction and rhythmic movement was consistent for all sites within the region of overlap even at the individual mouse level which was again assessed using correlation between group average and site level trajectories (**Table S1**).

Extending this analysis to encompass all four limbs and the mouth reveals that individual limb control is not spatially segregated on the cortex (**Fig. 5A**). Rather, complex motor movements (*i.e.* actions) exhibit topographical organization, emerging from synchronized and coordinated activities that reside within the overlap of cortical representations of individual limbs. For instance, the simultaneous activation of right and left forelimbs results in bilateral reaching motions, often coinciding with mouth opening, resembling an eating-like action, and primarily localized in M1 (**Fig. 5B**, Top). Similarly, the coordinated retraction of right and left forelimbs overlaps with hindlimb flexion, culminating in locomotion-like actions predominantly mapped in posterior somatosensory cortex, extending into parietal and retrosplenial regions (**Fig. 5B**, Bottom). These findings suggest the presence of cortical motor programs facilitating coordinated, multi-limbed movements reminiscent of natural behaviors.

## Discussion

Combining whole-hemisphere optogenetic stimulation with DLC and 3-dimensional motion capture across four camera views, we developed MOMM to simultaneously map motor output of the forelimbs, hindlimbs, and mouth in awake, head-fixed mice. MOMM facilitated the mapping of cortical representations of individual limb displacements in addition to finer articulations of specific limbs. Expanding the photostimulation grid across the entire left hemisphere revealed multiple cortical regions activating motor responses beyond the anatomical boundaries of somatomotor cortex. Additionally, we identified cortical sites that co-activated multiple limbs, producing coordinated movements resembling natural behaviors such as eating and locomotion. Overall, these findings highlight the topographical organization of cortical motor programs responsible for coordinated, multi-limbed, and behaviorally-relevant movements.

To our knowledge, MOMM is the first toolset capable of characterizing 3-dimensional motor movements of multiple limbs concurrently in awake mice without the use bulky electronics that hamper natural movement, or manual labeling of specific features, any of which preclude simultaneous mapping of multiple limbs. Critical to this methodology was extensive behavioral acclimation to the motor mapping system. Due to the difficulty in acclimation, most motor mapping in rodents thus far has been performed using anesthetized protocols which can suppress naturalistic movements ([39], [40], [20], [38], [41], [10], [11], [22]). Studies that have performed optogenetic mapping in awake mice have limited their scope to examining forelimb movements [21]. Our extensive and effective training paradigm successfully habituated mice to relax in a natural posture with both forelimbs and hindlimbs freely suspended, ensuring high fidelity movement tracking, and avoiding potential confounds of anesthesia on motor output. For all mapped body parts, photo-evoked movements exhibited high signal-to-noise ratio, were highly repeatable at the single trial level, and movement displacement was consistent within and across mice (**Fig. S1**).

Having established the fidelity of MOMM for tracking multiple features in 3D, we first validated the technique by mapping the well characterized representations of the forelimb. Within right forelimb, we see the spatial segregation of both rostral and caudal movements in line with several prior reports ([20], [10], [21]). The high fidelity of the motion tracking allows for dissecting finer forepaw movements, revealing that over M1, the right forepaw undergoes extension coupled with supination, resulting in a grasping-like motion. This presence of both gross translations and grasping motions can be attributed to distinct neural pathways: the C8-projecting layer 5b neurons control the distal forelimb (wrists and digits) involved in grasping, while C4-projecting corticospinal neurons (CSNs) govern proximal forelimb (shoulder and neck musculature) actions for gross translations [42]. The spatial superposition of these two movements highlights a potential integration within the motor control system. Intriguingly, these findings are replicated in the ipsilateral left forepaw, where photostimulation of the left hemisphere shows a comparable rostral-caudal organization. Further examination of finer articulations of the left forepaw also reveals a grasping-like motion, albeit with reduced magnitude. In mice, the corticospinal tract (CST) fibers originate in layer 5 pyramidal neurons, pass cross the midline at the pyramidal decussation, and descend in the contralateral dorsal column of the spinal cord [43]. Within the spinal cord, these fibers synapse onto interneurons, which subsequently modulate the activity of motor neurons [44]. However, about 10% of the CST axons in mice do not decussate at the pyramidal decussation, instead projecting to the ipsilateral spinal cord, thereby allowing the left hemisphere to innervate the left forelimb [44]. In addition, it is plausible that our stimulation not only activates corticospinal neurons but also engages local intrinsic circuits, some of which may include representations from the contralateral side. This ipsilateral innervation is supported by the strong underlying interhemispheric functional connectivity seen within both motor and somatosensory cortices ([30], [45], [46], [47], [48], [31]). Furthermore, photostimulation of left hemisphere sensory hindlimb (HLS1) results in activation of homotopic HLS1 in the right hemisphere [28]. However, despite the more precise control afforded by ChR2, there remains some uncertainty about the specific circuits involved or the directionality of innervation due to the contributions of either antidromic (axon to soma/dendrite) or orthodromic (dendrite/soma to axon) activity ([49], [50], [51]).

Mapping over the entire left hemisphere revealed that 64% of the mouse cortex is engaged in motor movement generation. Significant overlap among gross translation maps prompted an investigation into movement evoked by shared stimulation sites (**Fig. 4**), facilitated by MOMM’s simultaneous and high-fidelity tracking capabilities. Analysis of these overlapping regions identified time-locked bilateral forelimb movements elicited from both M1 (forelimb reach) and S1 (forelimb retract) (**Fig. 5A**). These findings are consistent with earlier studies ([52], [53], [11]) showing that stimulating the somatosensory cortex in rodents induces movement. In vivo imaging during a skilled forelimb retrieval task uncovered a logic for parallel-ordered corticospinal circuits, where specific subgroups of corticospinal neurons (CSNs) are activated in distinct cortical areas and sequences [54]. These circuits facilitate precise movements by preferentially connecting to spinal cord premotor neurons and their associated muscle groups [54]. This phenomenon may partially account for the extensive similarity observed across limb translation maps – specific CSN subgroups may drive individual limb translations, collectively contributing to the generation of complex actions.

Mapping multiple limbs over the cortex also revealed that motor movement representations extend beyond traditional somatomotor cortex boundaries, encompassing portions of the parietal and retrosplenial cortex. Specifically, 14% of stimulation sites evoking movement were located outside of typical somatomotor areas. Eating-like actions predominantly engage the somatomotor cortex, while locomotion-like actions are mainly associated with the parietal cortex, extending into the retrosplenial cortex. This localization aligns with LT-ICMS studies across various mammals including rats [11], rhesus monkeys [55], capuchin monkeys, [7], and macaque monkeys ([56], [57], [58]) that movements can be elicited from somatosensory and posterior parietal cortex stimulation. This localization also suggests the existence of a motor pathway within the parietal cortex, possibly integrated with or independent of the motor cortex.

Recent research in macaques has proposed two parallel networks in motor control—one involving M1 and another relatively independent [13]. Thus, regions of the parietal cortex may harbor a parallel corticospinal pathway capable of functioning somewhat autonomously from the motor cortex, potentially contributing to the locomotion-like actions observed within this region. Moreover, stimulation of different brain regions may elicit actions with distinct functional roles. For example, studies on whisking have demonstrated that M1 orchestrates rhythmic whisker protraction, whereas S1 governs short-latency retraction [59]. These distinct movements are mediated by two parallel pathways—a M1-brainstem pathway for rhythmic protraction and a S1-brainstem pathway for retraction [60]. The divergence between eating-like and locomotion-like actions in these regions underscores the notion that the former actions contribute to object manipulation and exploratory behaviors, while the latter actions are associated with avoidance behaviors.

The presence of these two distinct ethologically-relevant motor programs supports the emerging concept of an ethological organization principle [14] and that the functional organization of the cortex may be movement-based, rather than muscle based [10]. Studies in mice have shown that direct- and indirect-pathway spiny projection neurons (SPNs), found predominantly in the striatum, show locally biased spatiotemporal organization, with action-specific SPNs spatially clustered together [61]. Interestingly, different actions were not encoded by discrete SPNs but rather by overlapping SPN ensembles, with more similar actions encoded by closer ensemble representations (Klaus et al., 2017). This suggests that distinct SPN ensembles could underlie the eating-like and locomotion-like actions revealed through MOMM. Additionally, bidirectional connectivity of brain structures throughout the motor system (*i.e.* ascending and descending projections) allows for the function of a brain region to change during the behavioral “timescale” (acquisition, execution, or adaption) [62]. For example, the basal ganglia and cortex may contribute differently to skill acquisition but work in concert during its execution ([63], [64]). The topographical patterns observed in our maps may reflect our optogenetic stimulation actuating one aspect of several phases required to execute behavior, implying potential changes in these maps across different behavioral phases. While data-driven analysis of overlapping contours predominantly identified these two primary actions, empirical examination of photoevoked movements revealed further subdivisions, including discrete and rhythmic forelimb and hindlimb movements (**Fig. 3**). Employing machine learning to classify these actions without preconceived movement biases could offer a more nuanced action map. Additionally, exploring paired spatiotemporal dynamics across limbs would further validate the coordinated nature of multi-limb trajectories suggested by time-locked evolution of the multi-limbed trajectories.

In summary, we developed a Multilimb Optogenetic Motor Mapping (MOMM) method for examining cortical organization and execution of multi-limbed movements in awake mice. We identified the presence of cortical regions that encode motor programs for coordinated, multi-limbed, actions. We hope to further advance the utility of MOMM by exploring the neural correlates of coordinated movements through concurrent neuroimaging and motor mapping. MOMM opens the window to the application of awake motor mapping in determining the evolving relationships between motor circuit repair and motor recovery following injury.

## Author Contributions

**Nischal Khanal:** Data acquisition, software development, analysis, writing (original draft, review, and editing). **Jonah Padawer-Curry:** Experimental and system design, software development analysis, writing (original draft, review, and editing)**, Trevor Voss:** project conceptualization, experimental design, pilot data collection and analysis, **Kevin Schulte:** Experimental design, software development**, Brock Burke:** software development**, Annie Bice:** animal protocol, animal preparation, **Adam Bauer:** Project conceptualization, project administration, funding acquisition, supervision, Writing (review and editing).

## Acknowledgements

This work was supported by National Institute of Health grants R01NS126326 (AQB) R01NS102870 (AQB), RF1AG07950301 (AQB), Imaging Science Pathway T32 Grant T32EB014855 (NK, JPC). We want to thank Broc Burke for introducing us to DeepLabCut, Anmol Jarang and Evan Morris for help with designing and 3D printing the motor mapping mouse stand, and Sung Min Park for providing mouse illustrations for the figures. We also want to thank Jin-Moo Lee for provided valuable advice and interpretation of results.

## References

1. Penfield, W. and E. Boldrey, Somatic motor and sensory representation in the cerebral cortex of man as studied by electrical stimulation. Brain, 1937. 60(4): p. 389–443.

2. Penfield, W. and T. Rasmussen, The cerebral cortex of man; a clinical study of localization of function. 1950.

3. Nudo, R.J. and G.W. Milliken, Reorganization of movement representations in primary motor cortex following focal ischemic infarcts in adult squirrel monkeys. J Neurophysiol, 1996. 75(5): p. 2144–9.

4. Graziano, M.S., C.S. Taylor, and T. Moore, Complex movements evoked by microstimulation of precentral cortex. Neuron, 2002. 34(5): p. 841–51.

5. Griffin, D.M., et al., EMG activation patterns associated with high frequency, long-duration intracortical microstimulation of primary motor cortex. J Neurosci, 2014. 34(5): p. 1647–56.

6. Cooke, D.F., et al., Reversible Deactivation of Motor Cortex Reveals Functional Connectivity with Posterior Parietal Cortex in the Prosimian Galago (Otolemur garnettii). J Neurosci, 2015. 35(42): p. 14406–22.

7. Baldwin, M.K.L., et al., Representations of Fine Digit Movements in Posterior and Anterior Parietal Cortex Revealed Using Long-Train Intracortical Microstimulation in Macaque Monkeys. Cereb Cortex, 2018. 28(12): p. 4244–4263.

8. Cooke, D.F., et al., The functional organization and cortical connections of motor cortex in squirrels. Cereb Cortex, 2012. 22(9): p. 1959–78.

9. Baldwin, M.K., D.F. Cooke, and L. Krubitzer, Intracortical Microstimulation Maps of Motor, Somatosensory, and Posterior Parietal Cortex in Tree Shrews (Tupaia belangeri) Reveal Complex Movement Representations. Cereb Cortex, 2017. 27(2): p. 1439–1456.

10. Brown, A.R. and G.C. Teskey, Motor cortex is functionally organized as a set of spatially distinct representations for complex movements. J Neurosci, 2014. 34(41): p. 13574–85.

11. Halley, A.C., et al., Distributed Motor Control of Limb Movements in Rat Motor and Somatosensory Cortex: The Sensorimotor Amalgam Revisited. Cereb Cortex, 2020. 30(12): p. 6296–6312.

12. Heiney, S.A., G.J. Wojaczynski, and J.F. Medina, Action-based organization of a cerebellar module specialized for predictive control of multiple body parts. Neuron, 2021. 109(18): p. 2981–2994 e5.

13. Bresee, C.S., et al., Reversible deactivation of motor cortex reveals that areas in parietal cortex are differentially dependent on motor cortex for the generation of movement. J Neurophysiol, 2024. 131(1): p. 106–123.

14. Graziano, M.S.A., Ethological Action Maps: A Paradigm Shift for the Motor Cortex. Trends Cogn Sci, 2016. 20(2): p. 121–132.

15. Gordon, E.M., et al., A somato-cognitive action network alternates with effector regions in motor cortex. Nature, 2023. 617(7960): p. 351-359.

16. Stepniewska, I., P.C. Fang, and J.H. Kaas, Microstimulation reveals specialized subregions for different complex movements in posterior parietal cortex of prosimian galagos. Proc Natl Acad Sci U S A, 2005. 102(13): p. 4878–83.

17. Overduin, S.A., et al., Microstimulation activates a handful of muscle synergies. Neuron, 2012. 76(6): p. 1071–7.

18. Desmurget, M., et al., Neural representations of ethologically relevant hand/mouth synergies in the human precentral gyrus. Proc Natl Acad Sci U S A, 2014. 111(15): p. 5718–22.

19. Budri, M., E. Lodi, and G. Franchi, Sensorimotor restriction affects complex movement topography and reachable space in the rat motor cortex. Front Syst Neurosci, 2014. 8: p. 231.

20. Harrison, T.C., O.G. Ayling, and T.H. Murphy, Distinct cortical circuit mechanisms for complex forelimb movement and motor map topography. Neuron, 2012. 74(2): p. 397–409.

21. Hira, R., et al., Distinct Functional Modules for Discrete and Rhythmic Forelimb Movements in the Mouse Motor Cortex. J Neurosci, 2015. 35(39): p. 13311–22.

22. Brown, A.R., et al., Complex forelimb movements and cortical topography evoked by intracortical microstimulation in male and female mice. Cereb Cortex, 2023. 33(5): p. 1866–1875.

23. Mathis, A., et al., DeepLabCut: markerless pose estimation of user-defined body parts with deep learning. Nat Neurosci, 2018. 21(9): p. 1281–1289.

24. Wang, H., et al., High-speed mapping of synaptic connectivity using photostimulation in Channelrhodopsin-2 transgenic mice. Proc Natl Acad Sci U S A, 2007. 104(19): p. 8143–8.

25. Lee, J., et al., Opposed hemodynamic responses following increased excitation and parvalbumin-based inhibition. J Cereb Blood Flow Metab, 2021. 41(4): p. 841–856.

26. Ayling, O.G., et al., Automated light-based mapping of motor cortex by photoactivation of channelrhodopsin-2 transgenic mice. Nat Methods, 2009. 6(3): p. 219–24.

27. Thomson, A.M. and C. Lamy, Functional maps of neocortical local circuitry. Front Neurosci, 2007. 1(1): p. 19–42.

28. Lim, D.H., et al., In vivo Large-Scale Cortical Mapping Using Channelrhodopsin-2 Stimulation in Transgenic Mice Reveals Asymmetric and Reciprocal Relationships between Cortical Areas. Front Neural Circuits, 2012. 6: p. 11.

29. Bauer, A.Q., et al., Effective Connectivity Measured Using Optogenetically Evoked Hemodynamic Signals Exhibits Topography Distinct from Resting State Functional Connectivity in the Mouse. Cereb Cortex, 2018. 28(1): p. 370–386.

30. White, B.R., et al., Imaging of functional connectivity in the mouse brain. PLoS One, 2011. 6(1): p. e16322.

31. Bice, A.R., et al., Homotopic contralesional excitation suppresses spontaneous circuit repair and global network reconnections following ischemic stroke. Elife, 2022. 11.

32. Albertson, A.J., et al., Normal aging in mice is associated with a global reduction in cortical spectral power and network-specific declines in functional connectivity. Neuroimage, 2022. 257: p. 119287.

33. Seitzman, B.A., et al., Functional network disorganization and cognitive decline following fractionated whole-brain radiation in mice. Geroscience, 2024. 46(1): p. 543–562.

34. Nath, T., et al., Using DeepLabCut for 3D markerless pose estimation across species and behaviors. Nat Protoc, 2019. 14(7): p. 2152–2176.

35. Jung, J., et al., The structural connectivity of higher order association cortices reflects human functional brain networks. Cortex, 2017. 97: p. 221–239.

36. Lyamzin, D. and A. Benucci, The mouse posterior parietal cortex: Anatomy and functions. Neurosci Res, 2019. 140: p. 14–22.

37. Czajkowski, R., et al., Encoding and storage of spatial information in the retrosplenial cortex. Proc Natl Acad Sci U S A, 2014. 111(23): p. 8661–6.

38. Bonazzi, L., et al., Complex movement topography and extrinsic space representation in the rat forelimb motor cortex as defined by long-duration intracortical microstimulation. J Neurosci, 2013. 33(5): p. 2097–107.

39. Ramanathan, D., J.M. Conner, and M.H. Tuszynski, A form of motor cortical plasticity that correlates with recovery of function after brain injury. Proc Natl Acad Sci U S A, 2006. 103(30): p. 11370–5.

40. Hira, R., et al., Transcranial optogenetic stimulation for functional mapping of the motor cortex. J Neurosci Methods, 2009. 179(2): p. 258–63.

41. Silasi, G., et al., Improved methods for chronic light-based motor mapping in mice: automated movement tracking with accelerometers, and chronic EEG recording in a bilateral thin-skull preparation. Front Neural Circuits, 2013. 7: p. 123.

42. McKenna, J.E., G.T. Prusky, and I.Q. Whishaw, Cervical motoneuron topography reflects the proximodistal organization of muscles and movements of the rat forelimb: a retrograde carbocyanine dye analysis. J Comp Neurol, 2000. 419(3): p. 286–96.

43. Wang, Z., et al., Global Connectivity and Function of Descending Spinal Input Revealed by 3D Microscopy and Retrograde Transduction. J Neurosci, 2018. 38(49): p. 10566–10581.

44. Welniarz, Q., I. Dusart, and E. Roze, The corticospinal tract: Evolution, development, and human disorders. Dev Neurobiol, 2017. 77(7): p. 810–829.

45. Jonckers, E., et al., Functional connectivity fMRI of the rodent brain: comparison of functional connectivity networks in rat and mouse. PLoS One, 2011. 6(4): p. e18876.

46. Mechling, A.E., et al., Fine-grained mapping of mouse brain functional connectivity with resting-state fMRI. Neuroimage, 2014. 96: p. 203–15.

47. Grandjean, J., et al., Structural Basis of Large-Scale Functional Connectivity in the Mouse. J Neurosci, 2017. 37(34): p. 8092–8101.

48. Rahn, R.M., et al., Functional Connectivity of the Developing Mouse Cortex. Cereb Cortex, 2022. 32(8): p. 1755–1768.

49. Yizhar, O., et al., Optogenetics in neural systems. Neuron, 2011. 71(1): p. 9–34.

50. Lim, D.H., et al., Optogenetic approaches for functional mouse brain mapping. Front Neurosci, 2013. 7: p. 54.

51. Galvan, A., M.J. Caiola, and D.L. Albaugh, Advances in optogenetic and chemogenetic methods to study brain circuits in non-human primates. J Neural Transm (Vienna), 2018. 125(3): p. 547–563.

52. Neafsey, E.J., et al., The organization of the rat motor cortex: a microstimulation mapping study. Brain Res, 1986. 396(1): p. 77–96.

53. Tennant, K.A., et al., The organization of the forelimb representation of the C57BL/6 mouse motor cortex as defined by intracortical microstimulation and cytoarchitecture. Cereb Cortex, 2011. 21(4): p. 865–76.

54. Wang, X., et al., Deconstruction of Corticospinal Circuits for Goal-Directed Motor Skills. Cell, 2017. 171(2): p. 440–455 e14.

55. Rathelot, J.A., R.P. Dum, and P.L. Strick, Posterior parietal cortex contains a command apparatus for hand movements. Proc Natl Acad Sci U S A, 2017. 114(16): p. 4255–4260.

56. Cooke, D.F., et al., Complex movements evoked by microstimulation of the ventral intraparietal area. Proc Natl Acad Sci U S A, 2003. 100(10): p. 6163–8.

57. Gharbawie, O.A., et al., Multiple parietal-frontal pathways mediate grasping in macaque monkeys. J Neurosci, 2011. 31(32): p. 11660–77.

58. Mayer, A., et al., The Multiple Representations of Complex Digit Movements in Primary Motor Cortex Form the Building Blocks for Complex Grip Types in Capuchin Monkeys. J Neurosci, 2019. 39(34): p. 6684–6695.

59. Matyas, F., et al., Motor control by sensory cortex. Science, 2010. 330(6008): p. 1240-3.

60. Sreenivasan, V., et al., Parallel pathways from motor and somatosensory cortex for controlling whisker movements in mice. Eur J Neurosci, 2015. 41(3): p. 354–67.

61. Klaus, A., et al., The Spatiotemporal Organization of the Striatum Encodes Action Space. Neuron, 2017. 95(5): p. 1171–1180 e7.

62. Gmaz, J.M., et al., Integrating across behaviors and timescales to understand the neural control of movement. Curr Opin Neurobiol, 2024. 85: p. 102843.

63. Park, J., L.T. Coddington, and J.T. Dudman, Basal Ganglia Circuits for Action Specification. Annu Rev Neurosci, 2020. 43: p. 485–507.

64. Park, J., et al., Conjoint specification of action by neocortex and striatum. bioRxiv, 2023: p. 2023.10.04.560957.

