## Supplemental Figures for "Concurrent optogenetic motor mapping of multiple limbs in awake mice reveals cortical organization of coordinated movements"

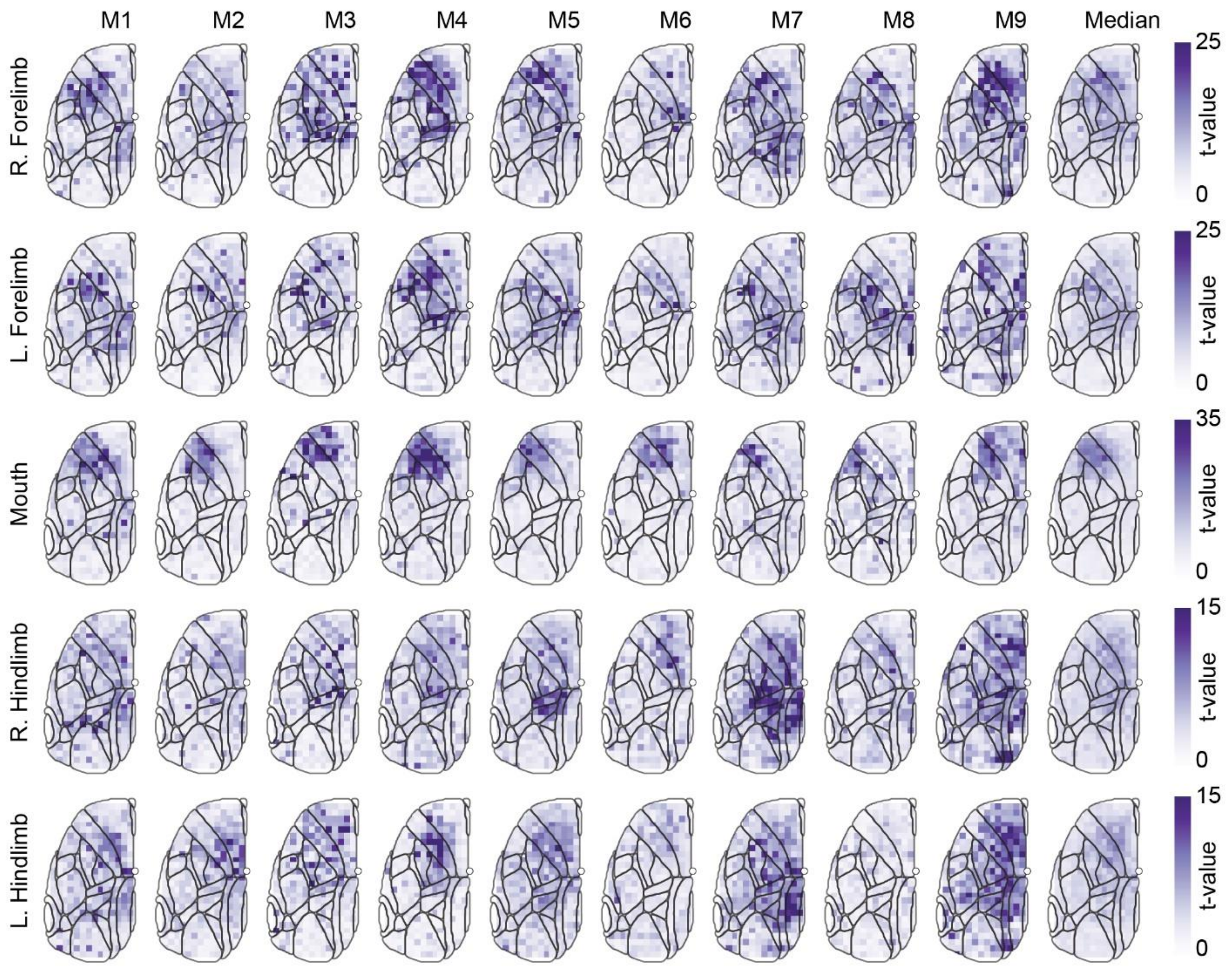


**Supplemental 1: Total Displacement – Mouse Level Variability**

The AUC displacement quantification at each site across all runs of each mouse was used to assess (via t-test, 5% significance level) the significance of the evoked movements (against the null of no movement). This metric was repeated for all four limbs and the mouth. The median map is generated using the median t-statistic value at each site across all nine mice. Higher t-values indicate that a movement generated at a specific site is more significant and this is reflected in that the median maps closely resemble the topographies of the gross translation maps (**Fig. 1E**).

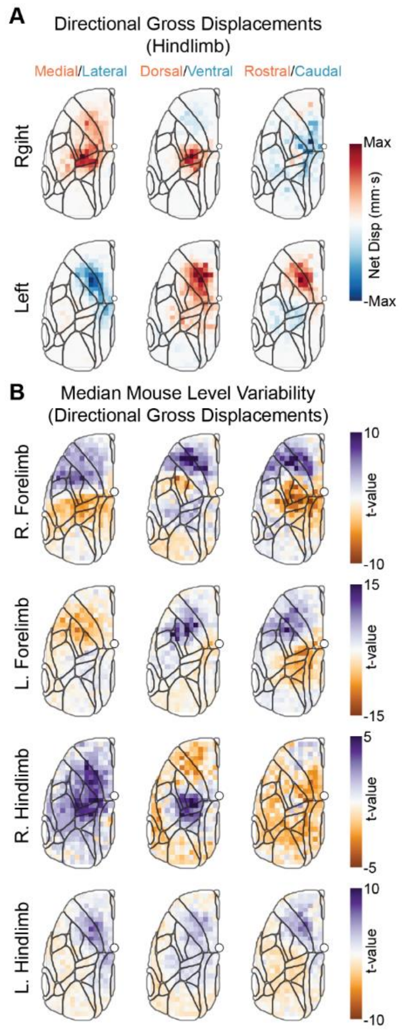


**Supplemental 2: Hindlimb directional displacements and directional displacement variability.**

A) Motor maps showing the directionality of right (top) and left (bottom) hindlimb movements along three directions. Positive (red) values indicate movement in the medial, dorsal or rostral directions while negative (blue) values indicate movement in the lateral, ventral or caudal directions. Maximum displacements of right hindlimb: 2.02 mm•s (M/L), 2.0 mm•s (D/V), 1.18 mm•s (R/C), left hindlimb: 1.93 mm•s (M/L), 0.89 mm•s (D/V), 1.83 mm•s (R/C). B) As in the gross translation case (**Fig. S1**), the AUC displacement quantification at each site across all runs of each mouse was used to assess the significance of the evoked movements. The median map displayed is generated using the median t-statistic value at each site across all nine mice.

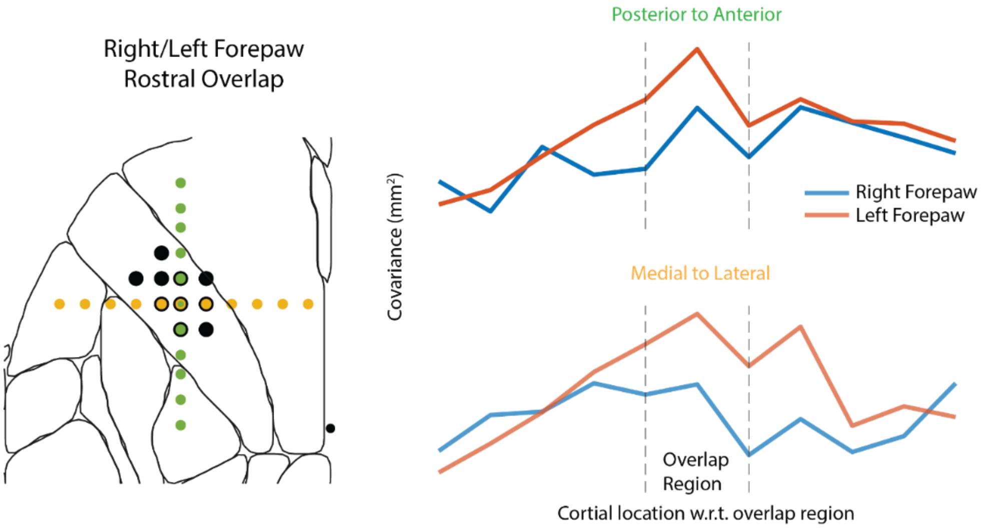


**Supplemental 3 – Specificity of trajectories**

The movement characteristics at sites surrounding the overlap region (black dots) for the primary representations of the two forelimbs was examined to quantify the spatial specificity of the co-activated movements. Sites extended anterior to posterior (green dots) and lateral to medial (yellow dots) through the region of overlap. The covariance between the group averaged trajectory at a given site and the group averaged trajectory over the region of overlap was computed, with large covariance values indicating more similar movements. The plots on the right show that the movements within the region of overlap more closely resemble the co-activated trajectory than the sites outside the region. Covariance values: Posterior RF: -0.793±10.0, Peak RF within overlap: 36.9, Anterior RF: 25.0±13.8, Posterior LF: 11.9±15.1, Peak LF within overlap: 55.1, Anterior LF:19.1±11.5. Medial RF: 20.6±12.2, Peak RF within overlap: 36.9, Lateral RF:19.5±9.90, Medial LF: 15.0±14.9, Peak LF within overlap: 55.1, Lateral LF:26.7±17.5.

**Table S1. Mouse Level Trajectory Correlation with Group Level Trajectory**

| Mouse | R/L Forelimb (RF) Correlation | R/L Forelimb (LF) Correlation | L Forelimb/R Hindlimb (LF) Correlation | L Forelimb/R Hindlimb (RH) Correlation |
|---|---|---|---|---|
| 1 | 0.87±0.072 | 0.82±0.10 | 0.64±0.16 | 0.92±0.039 |
| 2 | 0.92±0.051 | 0.81±0.14 | 0.86±0.096 | 0.86±0.029 |
| 3 | 0.87±0.10 | 0.84±0.086 | 0.92±0.040 | 0.87±0.075 |
| 4 | 0.94±0.015 | 0.77±0.088 | 0.92±0.047 | 0.90±0.039 |
| 5 | 0.96±0.020 | 0.89±0.065 | 0.94±0.028 | 0.92±0.029 |
| 6 | 0.89±0.059 | 0.78±0.11 | 0.77±0.073 | 0.96±0.024 |
| 7 | 0.92±0.014 | 0.77±0.15 | 0.80±0.14 | 0.87±0.035 |
| 8 | 0.96±0.016 | 0.82±0.096 | 0.92±0.037 | 0.88±0.087 |
| 9 | 0.91±0.11 | 0.86±0.098 | 0.89±0.052 | 0.94±0.020 |
